# Quantal release through the fusion pore

**DOI:** 10.64898/2026.11.23.768746

**Authors:** Alejandro Martínez-Valencia, Guillermo Ramírez-Santiago, Francisco F. De-Miguel

## Abstract

Exocytosis of neurotransmitters and peptides begins with the formation of a fusion pore that connects the vesicle lumen to the extracellular space. Release is completed when vesicles undergo full fusion. Here we show that full quanta of serotonin are released through a slowly closing fusion pore without full vesicle fusion. Amperometric recordings of serotonin release from dense-core vesicles in the axons of isolated Retzius neurons of the leech contained a subset of small release currents with a long biexponential decay. These slow-closure currents reflected the release of 90,262 ± 8,829 molecules, a number similar to full-fusion quanta and 54% larger than that of kiss-and-run events. The reconstructed dynamics of fusion pores show that slow-closure pores narrow rapidly after the peak and then close slowly or remain open. By contrast, kiss-and-run pores closed while vesicles still contained a large fraction of their serotonin content. Forces influencing pore kinetics were analyzed from pore velocity. The velocity decay of slow-closure pores developed a rapid undershoot characteristic of near-critical or underdamped relaxation. By contrast, in kiss-and-run pores, velocity decayed exponentially and turned negative during closure, consistent with an overdamped relaxation. Analysis of pore velocity revealed that slow-closure pores initiate rapid closure under a stiff force coupled to the pore diameter. Another force then arrests closure at a residual diameter of 1–2 nm that membrane tension alone cannot sustain. This allows release of the full serotonin quantum.

**Impact statement:** Release of the entire transmitter content of a vesicle, the quantum, is usually associated with the opening of a fusion pore followed by full vesicle fusion. Here we show that a full serotonin quantum can be released without full fusion, through pores that close rapidly to a residual diameter and then close slowly or stay open.

## Introduction

Neurons and endocrine cells release quantal pulses of signaling molecules when intracellular vesicles fuse with the plasma membrane (for review, see Katz, 1996; Hökfelt et al., 2018). Quantal release is evoked by a series of molecular processes. Elevations in the intracellular calcium level activate the tightening of the SNARE complex that links vesicles to the plasma membrane (for review see Sudhof, 2013). As a result, a nanometer-sized pore opens within a fraction of a millisecond and gradually expands (Heuser et al., 1979; Chandler and Heuser, 1980; Breckenridge and Almers, 1987a; Zimmerberg et al., 1987; Neher 1993). The nascent pore allows liquid to flow into the vesicle, hydrates the intravesicular matrix, and facilitates the efflux of soluble transmitter to the extracellular space (Chow et al., 1992; Wightman et al., 1995; Staal et al., 2004; Amatore et al., 2005). The pore expansion is usually followed by full vesicle fusion and extrusion of the quantum remains. In some cases, the pore opens and then closes (Breckenridge and Almers, 1987b; Wightman, et al, 1991; Alvarez de Toledo et al., 1993). Such phenomenon, known as “kiss and run” (Ceccarelli et al., 1973; Alabi and Tsien, 2013) allows only a fraction of the quantum to be released (MacDonald et al., 2006). Here we show that whole quanta of 5-HT can be released through pores that open and close slowly or remain open without occurrence of full vesicle fusion.

Our experiments involved electrochemical recordings of 5-HT release from the large axon of the classical serotonergic Retzius neuron of the leech. Retzius neurons in culture release small serotonin (5-HT) quanta containing ∼4,000 molecules from clear synaptic vesicles, and much larger quanta with >90,000 molecules from extrasynaptic dense core vesicles in the soma, axon and perisynaptic regions (Kuffler et al., 1987; Bruns and Jahn, 1995; Bruns et al., 2000; Trueta et al., 2003). The amplitude and kinetics of amperometric currents recorded upon 5-HT oxidation allow a clear distinction between individual releases from either vesicle type (Bruns and Jahn, 1995).

The present report is based on the incidental observation that amperometric recordings of 5-HT exocytosis from axonal dense-core vesicles display small events, similar to kiss-and-run but with prolonged relaxations in the absence of a fusion spike. To test the hypothesis that entire quanta are released through pores that remain open for extended periods, we quantified serotonin release from full fusion, kiss-and-run, and slow-closure events. Then the pore diameter for each event was reconstructed using a toroidal pore geometry (De-Miguel, 2026a). The forces governing opening and closure of the pore were estimated from the velocity of its diameter.

## Results

### Modes of 5-HT exocytosis

Gentle pressure applied to the axon membrane with the tip of the amperometric electrode evoked bursts of exocytosis from 5-HT-filled dense-core vesicles. Individual exocytotic events resembled those induced by ionomycin, which increases the intracellular calcium levels (Bruns et al., 2000).

The exocytic currents fell into three kinetic types (Figure 1). Most events (85%) were the well-known large, rapid spikes produced by full vesicle fusion (full-fusion currents). In 17% of events, the spike was preceded by a foot reflecting 5-HT efflux through the expanding pore (Figure 1A). The 5-HT in the foot and the spike together formed the quantum. A second type (13% of events) comprised kiss- and-run currents, which rose rapidly and decayed as a single exponential (R² > 0.97), because the pore opened and closed without full fusion (De-Miguel, 2026b). The third type (10% of events), which prompted this report, comprised slow-closure currents. They were smaller than full-fusion spikes and decayed biexponentially, with τ₁ = 1.9 ± 0.2 ms and τ₂ = 17.5 ± 2.4 ms (R² > 0.98; Figure 1A–C; Table 1). The recorded currents, analyzed without fitting, confirmed that the two types differ in their late decay (Figure 1—figure supplement 1).

**Figure 1.**
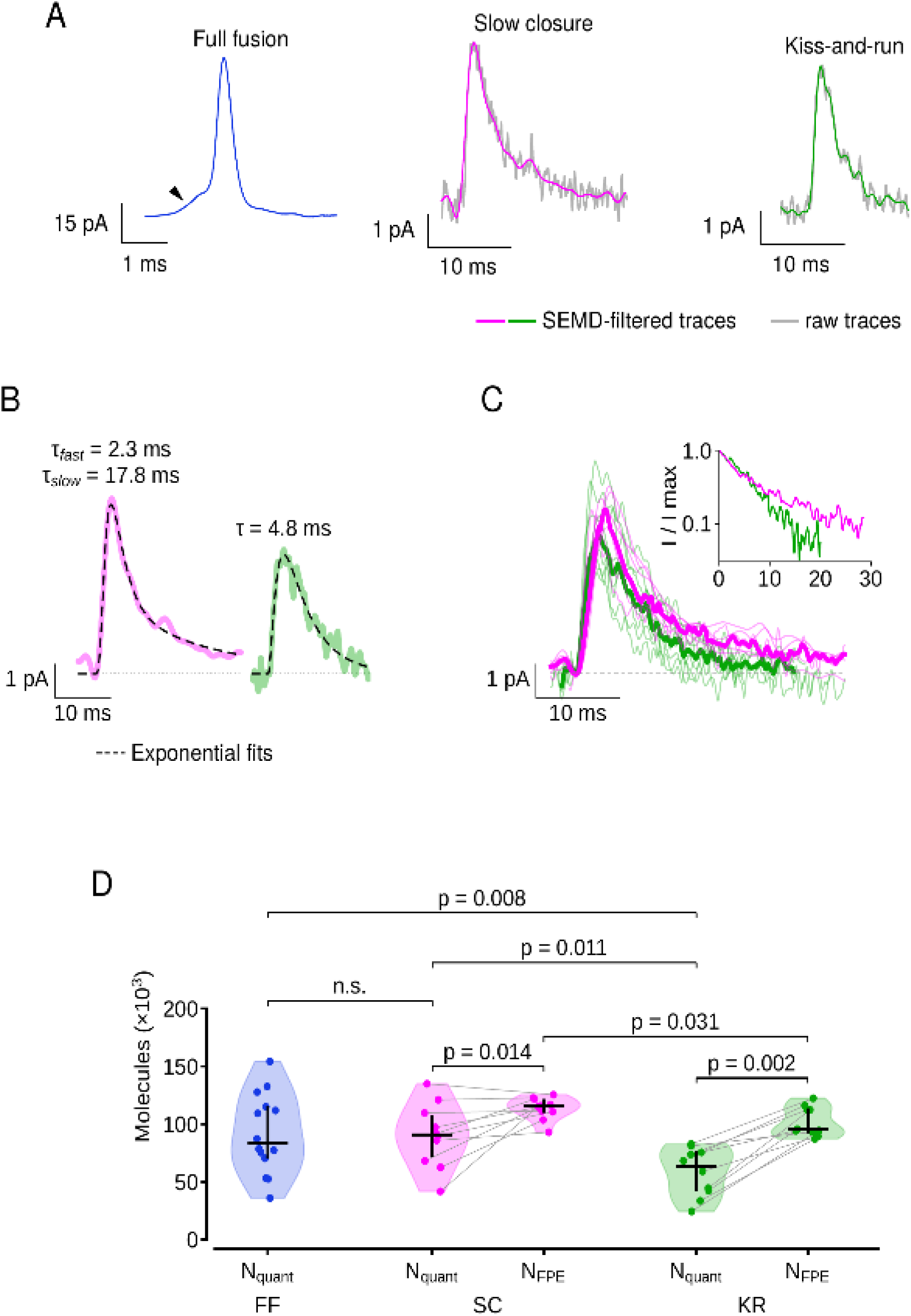
Three modes of 5-HT exocytosis from dense-core vesicles. A) Rapid full-fusion spike preceded by a foot (arrowhead, blue); small slow-closure current with biexponential decay (magenta); small kiss-and-run current with monoexponential decay (green). Note scale differences. SEMD-filtered traces are overlaid onto the original traces (gray). B) Median slow-closure and kiss-and-run events with fits of Equation 1 (dashed); the dotted line marks the pre-onset baseline. C) Slow-closure (n = 7) and kiss-and-run (n = 8) currents aligned at onset (thin lines); Medians are thick lines; the semilogarithmic inset shows the median decays normalized to peak values, highlighting biexponential and monoexponential decays. D) 5-HT molecules released (N_quant_) by full-fusion (FF, n = 14), slow-closure (SC, n = 10) and kiss-and-run (KR, n = 10) events; N_FPE_ is the vesicular content estimated by frozen pore extrapolation; gray lines join values of the same event. Violins show kernel density estimates, dots individual events, horizontal bars medians and vertical bars interquartile ranges. Welch’s t-test between groups and Wilcoxon signed-rank test (Wilcoxon, 1945) between N_quant_ and N_FPE_; n.s., not significant. A consistent color scheme is used across all figures.

**Table 1.** Parameter values for the pore release modes. Data are shown as mean ± SEM; rise exponents and damping ratios are given as median (range). Kinetic parameters were obtained by fitting Equation 1 to the fitted currents, and pore diameters were obtained from the toroidal reconstruction (Materials and methods). ‡Diameter and area at the end of the foot, reconstructed using the mean quantum of full-fusion events. §Quantum of full-fusion events (n = 14). Damping ratios are from eight slow-closure and eight kiss-and-run events (Materials and methods). Foot values exclude two foot currents removed from the rise analysis (Materials and methods).

| **Parameter** | **Slow-closure<br>(n=10)** | **Kiss and run<br>(n=10)** | **Foot currents<br>(n=16)** |
|---|---|---|---|
| Peak amplitude (pA) | 4.79 ± 0.29 | 3.96 ± 0.41 | 3.40 ± 0.61 |
| τ<sub>0</sub> (rise time constant, ms) | 2.13 ± 0.34 | 1.15 ± 0.16 | 3.07 ± 0.53 |
| Time to peak (ms) | 3.23 ± 0.25 | 2.84 ± 0.21 | 4.45 ± 0.40 |
| Rise exponent n, median (range) | 3.85 (2.0–4.48) | 2.0 (2.0–4.0) | 1.43 (0.73–2.51) |
| τ<sub>1</sub> (fast decay time constant, ms) | 1.9 ± 0.2 | 7.4 ± 1.5 | — |
| τ<sub>2</sub> (slow decay time constant, ms) | 17.5 ± 2.4 | — | — |
| Peak pore diameter (nm) | 2.82 ± 0.07 | 2.63 ± 0.12 | 2.56 ± 0.37‡ |
| Peak pore area (nm²) | 6.27 ± 0.34 | 5.54 ± 0.52 | 6.73 ± 1.95‡ |
| Molecules released | 90,262 ± 8,829 | 58,543 ± 6,626 | 91,585 ± 9,201§ |
| ζ Damping ratio ζ, median (range) | 1.13 (0.79–1.88) | 1.79 (0.98–3.63) | - |

### Molecules released by each release mode

The number of 5-HT molecules released in each event type is shown in Figure 1D. 5-HT molecules counted from charge (Nquant) are compared with vesicular content estimated by frozen pore extrapolation (N_FPE_; De-Miguel, 2026a; Materials and methods). Full-fusion events released 91,585 ± 9,201 molecules (mean ± SEM; n = 14), like the number reported by Bruns et al. (2000). Slow-closure events released a similar number with comparable variability, 90,262 ± 8,829 (n = 10; Welch’s t-test, p = 0.92), whereas kiss-and-run events released only 58,543 ± 6,626 molecules, accounting for 64% of the full-fusion amount (n = 10; p = 0.008). The N_FPE_ method predicts higher numbers of 5-HT molecules inside vesicles. Vesicles producing slow-closure currents contained 113,719 ± 3,068 molecules (Figure 1D), and kiss-and-run vesicles contained 101,831 ± 4,000 molecules (p = 0.32, compared to full fusion). Full-fusion counts are not possible with this method because the current peak remains undetermined. This result suggests that the quantum is marginally smaller than the number of molecules packed inside vesicles (References).

### Rise of amperometric currents

The early rise of amperometric currents is typically not considered in release analyses because it is masked by hydration of the intravesicular matrix and dissociation of the transmitter from it (Chow et al., 1992; Wightman et al., 1995; Amatore et al., 2005; Borges et al., 2010). However, each type of amperometric current could be clearly distinguished by two characteristics of its rising phase (Figure 2A–C), namely the n exponent that defined the onset delay and the τ₀ value. Slow-closure currents rose with the most delayed onset of the three types, with an exponent n∼4 (median; range 2.0–4.48) and a τ₀ of 2.13 ± 0.34 ms (median 1.75 ms; 1.27–4.71 ms). Kiss-and-run currents had a lower n=2.0 (2.0– 4.0; Mann–Whitney U test, p = 0.030) and a τ₀ about half that of slow-closure currents (1.15 ± 0.16 ms; median 1.01 ms; 0.68–2.39 ms; p = 0.006). Full-fusion feet had the lowest n=1.43 (0.73–2.51; p < 0.001 versus both slow-closure and kiss-and-run currents) and a τ₀ (3.07 ± 0.53 ms; median 2.25 ms; 1.0–6.91 ms) similar to that of slow-closure currents (p = 0.41) and longer than that of kiss-and-run currents (p = 0.001; Table 1).

**Figure 2.**
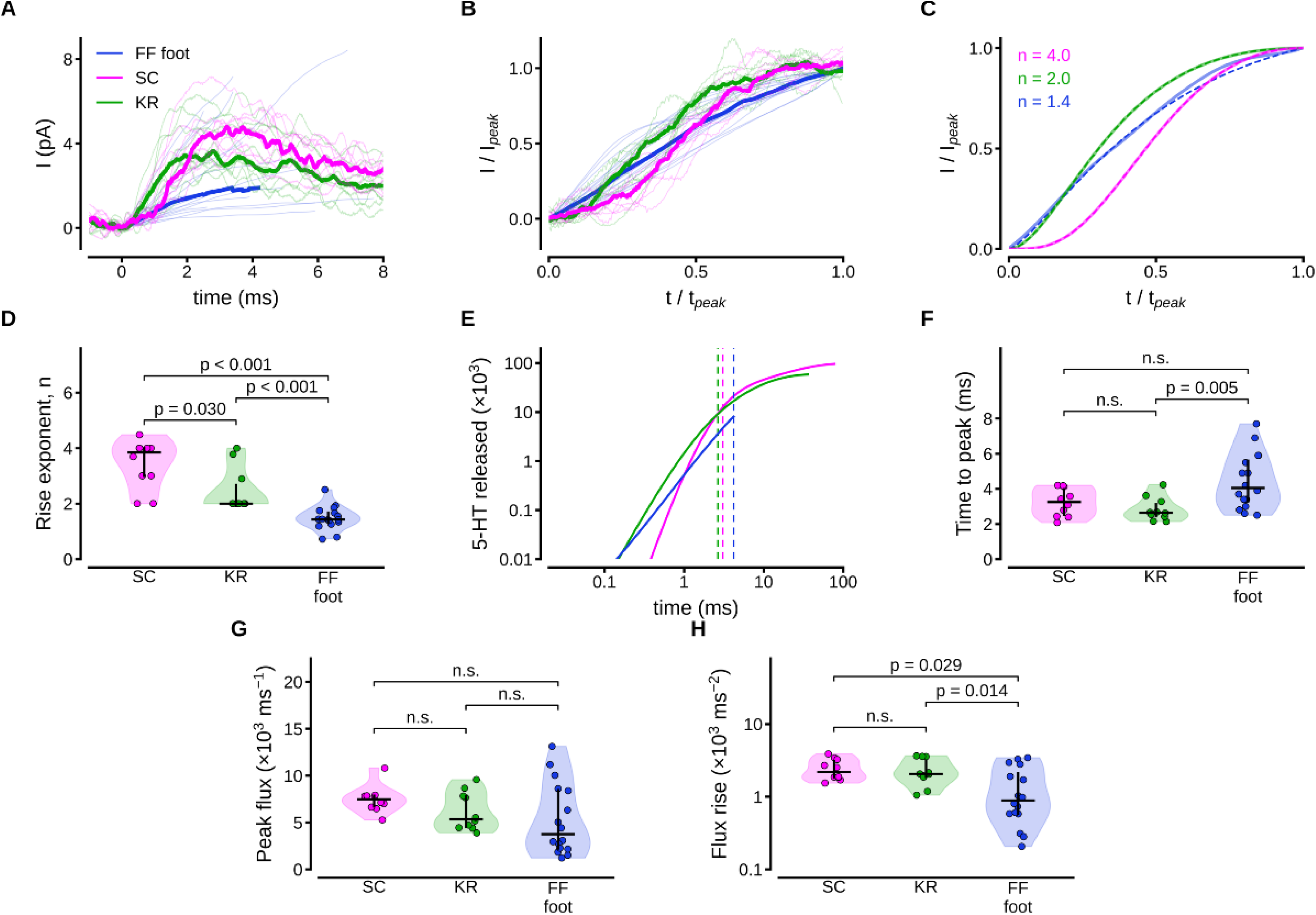
Release through the rise of amperometric currents. A) Rising phases of recorded currents aligned at onset (thin lines); thick lines are medians; n = 9 slow-closure, 10 kiss-and-run, and 16 foot currents. B) Rising phases normalized to peak amplitude and time to peak. C) Fits of median currents in B using Equation 1. Rise exponents n are indicated. D) Rise exponent for each event. E) 5-HT molecules released versus time (logarithmic). Slow-closure and kiss-and-run traces end at plateau; foot trace ends at spike onset. Vertical lines mark peak current times. The delay of the slow-closure trace reflects its sigmoidal onset. Early in the rise, the slow-closure current (n ≈ 4) releases fewer molecules than the kiss-and-run (n = 2) and foot (n ≈ 1.4) currents. Later, release rises more steeply. F) Time to peak. G) Peak flux. H) Rate of flux rise (logarithmic peak flux divided by time to peak). Violins show kernel density; horizontal bars are medians; vertical bars are interquartile ranges; Mann–Whitney U test; n.s. = not significant.

Slow-closure currents reached a maximum amplitude of 4.79 ± 0.29 pA and a peak flux of 7,476 ± 455 molecules per millisecond. This was statistically similar to kiss-and-run currents (3.96 ± 0.41 pA) and their peak flux (6,187 ± 644 molecules per millisecond; Mann–Whitney U test, p = 0.14 for both; Figure 2G). The foot preceding full fusion varied considerably in size, as in other cells (Jankowski et al., 1993), reaching 3.40 ± 0.61 pA and 5,309 ± 956 molecules per millisecond (n = 16), similar to slow-closure (p = 0.097) and kiss-and-run currents (p = 0.26; Figure 2G; Table 1).

### Currents predicted through frozen pores

To determine whether the current decay through slow-closure pores reflects two states of pore closure, recorded slow-closure currents were compared with currents reconstructed by freezing the pore diameter (De-Miguel, 2026a; Materials and Methods), first at the current peak and then at the inflection point 8 of the decay in slow-closure pores. Figure 3A shows that current through the pore frozen at its peak diameter decayed more slowly ( = 13.7 ± 0.5 ms) than the early decay of the recorded current (= 1.9 ± 0.2 ms; Wilcoxon signed-rank test, p = 0.002). Similar results across the slow-closure population suggest that the rapid post-peak current decay reflects pore closure (Figure 3A, inset). This result contrasts with an equivalent analysis of kiss-and-run currents (Figure 3B; De-Miguel, 2026b), in which the time constant of the current through the frozen pore ( = 15.7 ± 1.1 ms, range 10.6–22.0 ms) was twice that of the recorded current ( = 7.4 ± 1.5 ms, range 2.25 to 16.4 ms; p = 0.002; Table 1). Note that of frozen pore currents were similar regardless of the type of pore closure (Mann–Whitney U test, p = 0.19). When the pore was frozen right at the inflection between fast and slow closing (Figure 3C), the time constant of the slow decay current (τ_2_ = 17.5 ± 2.4 ms, range 4.75–28.2 ms) was similar (p = 0.064) to that of currents through frozen pores (τ= 26.2 ± 2.5 ms, range 15.1– 42.7 ms). Such similarity reflects the variance in the recorded currents (Figure 3C, inset).

**Figure 3.**
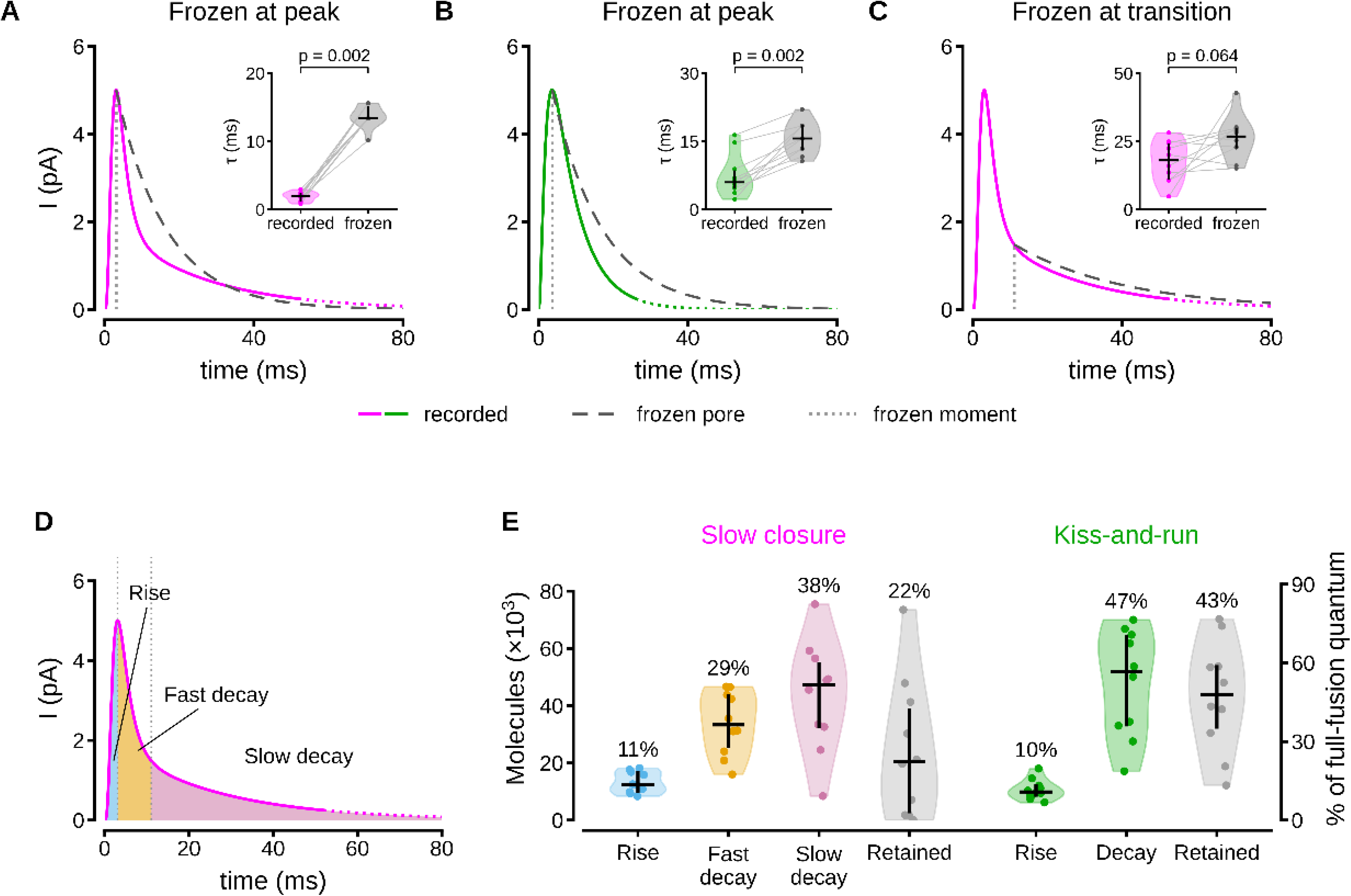
Release at different stages of pore dynamics. A) Median slow-closure current and frozen pore current at current peak. Inset: fast decay time constant (τ_1_) of recorded and frozen-pore currents (n = 10); lines join values of the same event. B) Kiss- and-run current and current through its pore frozen at current peak. Inset as in A. C) Slow-closure event in A with pore frozen at the transition between fast and slow decay components. Inset as in A. D) Phases of slow-closure events; kiss-and-run currents had only rise and decay. E) Molecules released in each phase and molecules retained in the vesicle (N_FPE_ minus molecules released; n = 10 each). Numbers above violins are mean percentages of the N_FPE_; Right axis, percentage of full-fusion quantum.

Figure 3D, E compares the percentage of 5-HT released in each phase of the slow-closure and kiss-and-run currents, relative to the N_FPE_ estimate of vesicle content. In slow-closure events, the fraction released increased from 11 ± 1% during the rise (14% of the full-fusion quantum) to 29 ± 3% during the fast decay (37% of the quantum) to 38 ± 5% during the slow decay (47% of the quantum). Kiss-and-run events released a similar fraction during the rise (10 ± 1%, 12% of the quantum; p = 0.31). Their single decay phase released 47 ± 6% (52% of the quantum). This is more than the fast decay of slow-closure currents (p = 0.031) but less than the fast and slow decays combined (68 ± 6%; p = 0.045). Conversely, the estimated fraction retained in the vesicle increased from 22 ± 7% in slow-closure events to 43 ± 6% in kiss-and-run events (p = 0.045).

### Pore dynamics

The diameter dynamics of both pore types reconstructed with toroidal geometry are shown in Figure 4. The first nanometer of the rising phase was excluded because the early current was contaminated by hydration of the intravesicular matrix (Chow et al., 1992; Wightman et al., 1995; Amatore et al., 2005; Borges et al., 2010). Capacitance recordings also show that the pore opens abruptly with a conductance of tens to a few hundred picosiemens, comparable to that of a large ion channel and corresponding to a diameter of about 1 nm (Breckenridge and Almers, 1987a; Spruce et al., 1990; Lollike et al., 1995; Albillos et al., 1997; Klyachko and Jackson, 2002; He et al., 2006). This is consistent with simulations in which pores narrower than about 1 nm are separated by an energy barrier from larger pores governed by the line tension of their edge (Tolpekina et al., 2004b; Wohlert et al., 2006).

**Figure 4.**
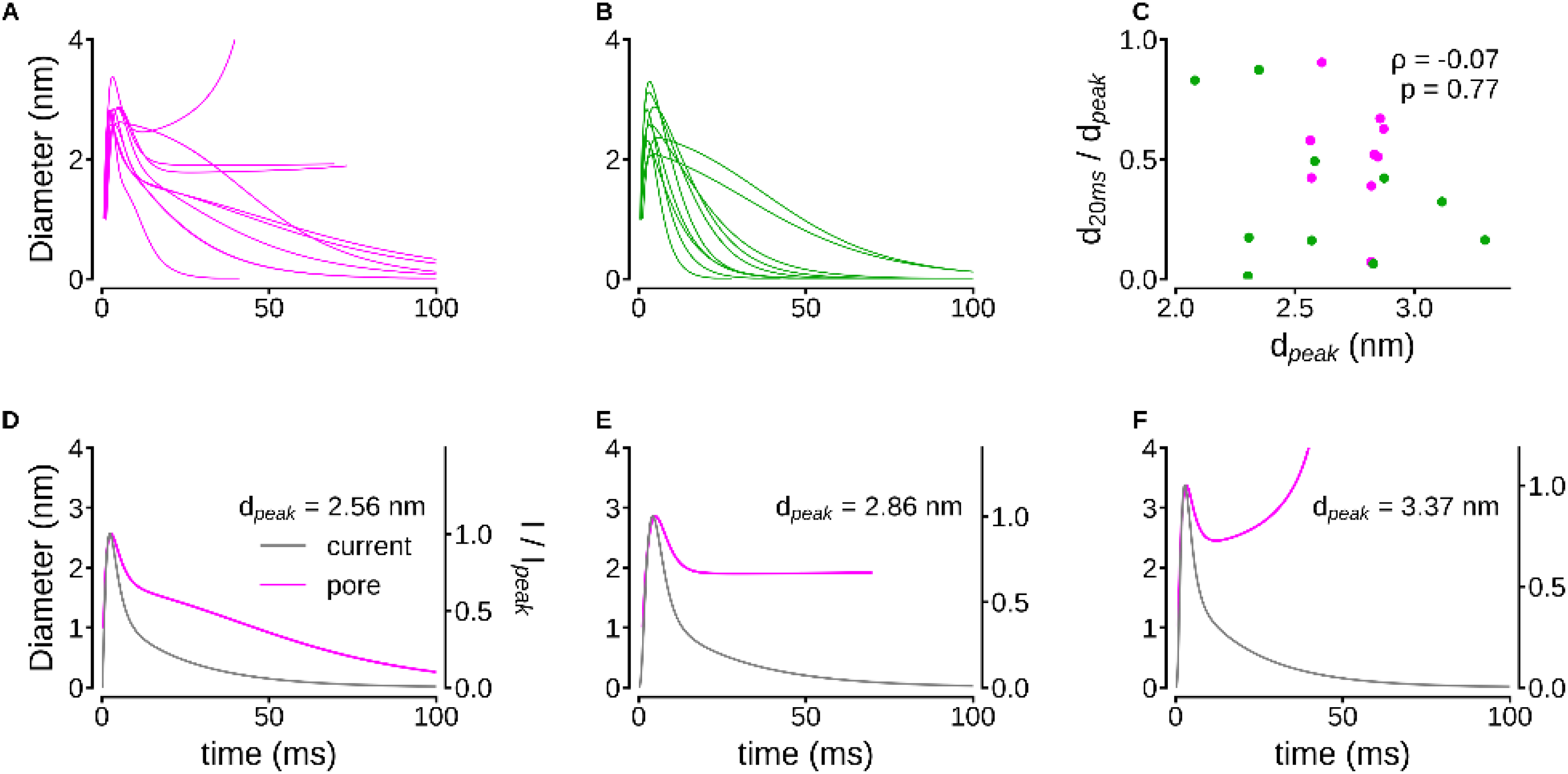
Reconstruction of pore diameter. A) Pore diameter for slow-closure events (n = 10). B) Pore diameter for kiss-and-run events (n = 10). C) Fraction of peak diameter remaining 20 ms after onset (d20 ms/dpeak) versus peak diameter (slow-closure pore F excluded). Spearman correlation. D–F) Pore diameter and normalized current for three slow-closure pores: D) fast narrowing followed by slow closure; E) a plateau at about 1.9 nm; F) late re-expansion in the largest pore, an artifact of an underestimated vesicle content (see text).

Reconstructed pores reached their peak diameter at 3.9 ± 0.4 ms in slow-closure events and 3.6 ± 0.4 ms in kiss-and-run (Mann–Whitney U test, p = 0.52). Peak diameters were 2.82 ± 0.07 nm (range 2.56–3.37 nm; n = 10) for slow-closure and 2.63 ± 0.12 nm (2.08–3.30 nm; n = 10) for kiss-and-run (p = 0.31). The highly variable pore producing the foot of full-fusion events reached 2.56 ± 0.37 nm (0.97–6.25 nm; n = 16), similar to either (p = 0.54 and p = 0.36).

After the peak slow-closure pores followed variable time courses (Figure 4A), from the diameter falling rapidly then declining more slowly (Figure 4D), to declining and then remaining open (Figure 4E), to falling and then reopening (Figure 4F). The late re-expansion is an artifact of an underestimated vesicle content: this event released more than its N_FPE_, and its reconstructed pore closes monotonically when the content is raised by 1%. These three slow-closure pores had peak diameters of 2.56, 2.86 and 3.37 nm. One pore developed a broad, weakly biexponential hump, rather than fast closure, resembling the decay of two kiss-and-run pores (Figure 4B; Figure 4—figure supplement 1A). Kiss-and-run pores closed monotonically after the peak (Figure 4B; De-Miguel, 2026b). Within 20 ms, their diameter had fallen to 35 ± 10% of the peak value.

Pore decay did not correlate with peak diameter in either the fast or slow phases of slow-closing pores (ρ = −0.13 and 0.07) or in monotonic kiss-and-run pores (Spearman ρ = −0.15, p = 0.68). Regardless of pore type, the fraction of peak diameter remaining after 20 ms was also unrelated to the maximum peak diameter (ρ = −0.07, p = 0.77; n = 19; Figure 4C). Pores that reached the same peak diameter, and therefore carried nearly the same peak 5-HT efflux, could close at very different rates (Figure 4—figure supplement 1B, C).

### Continuity between slow-closure and kiss-and-run events

Despite their diversity, slow-closure pores shared an exponential decay as a distinctive feature. In early stages, kiss-and-run and slow-closure pores were remarkably similar. Once wider than 1 nm, they opened at the same rate. They reached similar peak diameters (p = 0.31), similar peak fluxes (p = 0.14), and similar current amplitudes (p = 0.14). Moreover, they released similar numbers of molecules during the rise. These data suggest that the two pore types differed before the measurable pore: slow-closure currents rose with a larger exponent and a longer time constant (Table 1), placing the difference at the sub-nanometer stage of pore formation. Assuming that both pore types form from the same basic structure, their similarities and overlaps suggest a continuum, set by the strength of the forces acting on the pores from their early stage. To test this possibility, five properties of the rising phase (peak amplitude, time to peak, 0, rise exponent n, 12 and maximal rate of rise) and five properties of the decay (time from the peak to 50% and to 20% of the peak amplitude, fraction of the molecules released later than 10 ms after the peak, total number of molecules released, and share of the slow decay component) were analyzed by principal component analysis. The slow-component share and the number of molecules released were excluded, as they were used to classify the events. The two groups overlapped along their full range, with a unimodal joint distribution (Hartigan’s dip test, p = 0.50). Moreover, an unsupervised two-cluster partition failed to recover the two event types (Rand index 0.21), and the properties of the rising phase correctly classified the events in 70% of cases using leave-one-out logistic regression.

Within this continuum, the kinetics of the rising phase predicted the decay rate independently of event type. The time to peak correlated with the time to 50% decay (Spearman ρ = 0.63; within kiss-and-run events, 0.67; within slow-closure events, 0.73; p = 0.001 in a rank regression adjusted for event type, significant after Bonferroni correction for 25 comparisons). Faster rises were followed by faster decays, a pattern most evident in slow-closure events (ρ = −0.84, p = 0.002).

The first rising-phase component explained 36% of the variance in decay time, whereas event type explained only 2%. By contrast, the differences between the two event types in the slow component (late fraction of release and number of molecules released) were explained by event type rather than by the kinetics of the rising phase. These results reinforce the idea that graded forces acting on the early pore set the rate of subsequent relaxation in both event types. The transition to the slow phase that completes quantal release depends on an additional factor.

### Velocity of the pore

Having inferred that slow closure and kiss-and-run events belong to the same population responding to different forces, our next step was to estimate the forces 13 that determine the categories of pore closure. Because pores develop at the nanometer scale, where inertia is negligible, their dynamics reflect the balance between the forces acting on the pore and the friction opposing them, and therefore provide insight into the forces driving pore kinetics (Sandre et al., 1999; Brochard-Wyart et al., 2000; Chizmadzhev et al., 2000; Ryham et al., 2011). We first calculated the Reynolds number, Re = ρvL/η, to compare inertial and viscous forces acting on the pores. Using the density and viscosity of water (ρ = 10³ kg m⁻³; η = 10⁻³ Pa s), pore diameters L of 1–4.3 nm, and our measured velocities (0.04–12 nm/ms), we obtained Re ≈ 10⁻¹¹–10⁻⁷. In this low-Reynolds-number regime, viscous drag is linearly related to velocity, and the forces acting on the pore balance at every instant (Purcell, 1977; Happel and Brenner, 1983). Consequently, the pore velocity directly reflects the net driving force divided by an effective friction coefficient (Chizmadzhev et al., 2000).

Figure 5 compares diameter and velocity across both pore types. In flux-matched pairs, slow-closure pores showed a reproducible, sharp negative velocity undershoot, absent or much more gradual during kiss-and-run events (Figure 5). Aligned to the end of opening and normalized to peak negative velocity, slow-closure undershoots were narrower (Figure 5D). They reached minimum velocity earlier (median 2.0 ms, range 1.2–3.4 ms, vs. 3.6 ms, 2.6–8.1 ms; n = 8 each; Mann–Whitney U test, p = 0.015; Figure 5E), recovered to half-minimum faster (3.7 ms, 1.6–5.0 ms, vs. 9.6 ms, 4.1–19.2 ms; p = 0.001), and lasted half as long (half-width 5.5 ms, 2.5–7.4 ms, vs. 12.3 ms, 6.1–26.6 ms; p = 0.001; Figure 5F). The undershoot amplitude, however, was similar (−0.18 nm/ms, range −0.50 to −0.14 nm/ms, vs. −0.18 nm/ms, −0.33 to −0.09 nm/ms; p = 0.72), confirming a difference in kinetics rather than amplitude. In velocity–diameter plots (Figure 5— figure supplement 1), kiss-and-run trajectories moved along negative velocities directly to near-zero diameters, whereas slow-closure trajectories showed two negative phases: an initial undershoot returning to near-zero velocity at 1.5–2 nm, followed by slow constriction (Figure 5—figure supplement 1B).

**Figure 5.**
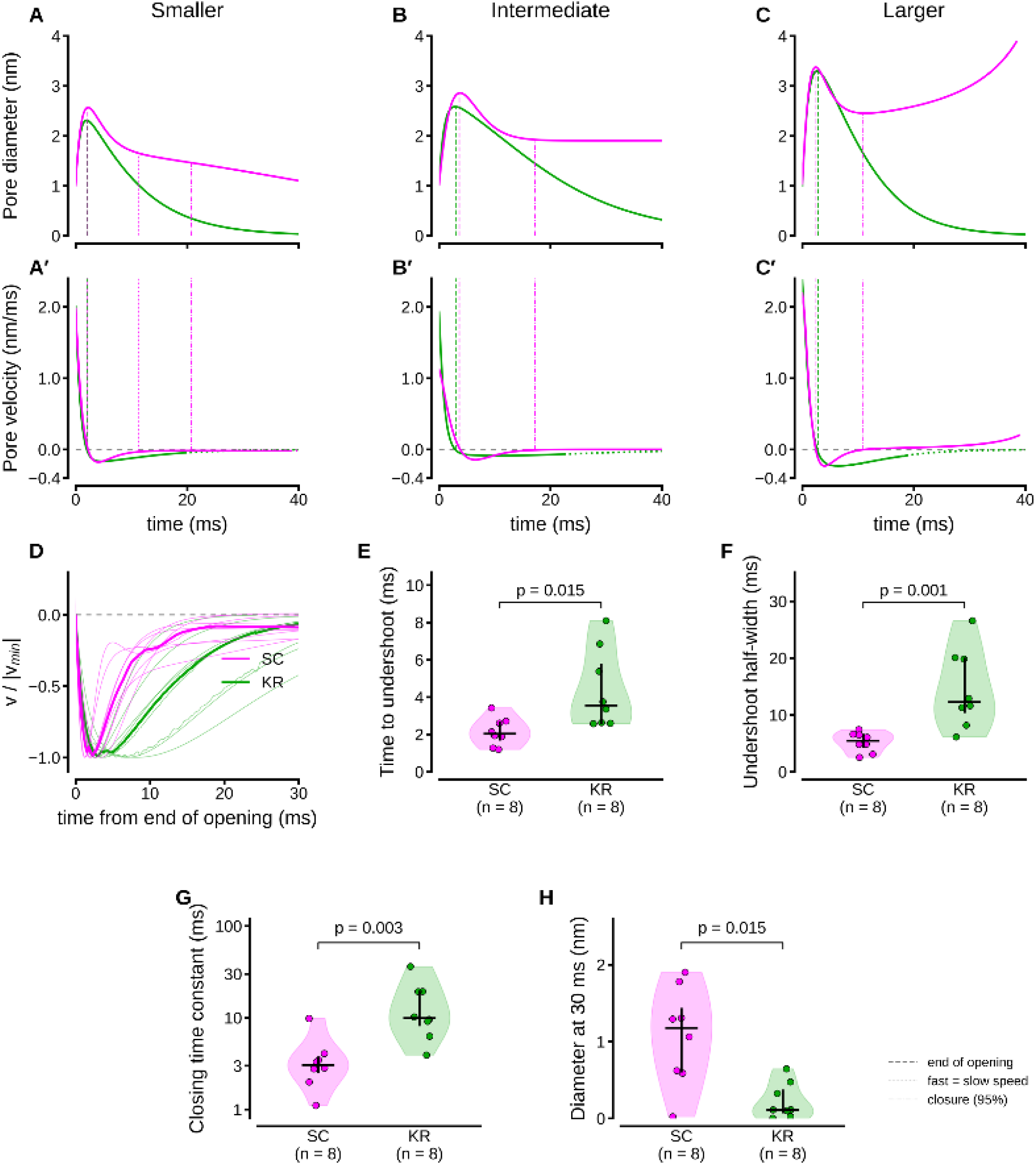
Pore diameter and velocity. A–C) Temporal development of pore diameter for slow-closure and kiss-and-run events with increasing amplitudes. Vertical lines mark the end of opening (diameter peak) and the crossing time between fast and slow closing phases (dotted), and 95% closure (dash-dotted only in A). A′–C′) Kinetics of pore velocity for the same pairs and vertical lines. D) Negative pore velocity during early closure, normalized to its minimum and aligned at the end of opening. Thin lines, individual events; thick lines, medians. E) Time from the end of opening to minimum velocity. F) Half-width of the velocity undershoot. G) Closing time constant (fast phase in pores with two closing phases); note the logarithmic scale. H) Pore diameter 30 ms after crossing 1 nm. In D–H, n = 8 per group, excluding pores that closed as a broad hump (Figure 4— figure supplement 1A) and the slow-closure event in C; violins show kernel density estimates, dots individual events, horizontal bars medians, and vertical bars interquartile ranges; Mann–Whitney U test. In C and C′, the closing phase was analyzed to the minimum diameter.

### Forces predicted by damped dynamics

The faster undershoot into negative velocities in slow-closure pores (Figure 5D) suggested that pore velocity could be modeled as a damped relaxation process characterized by an attenuation factor ζ. In this framework, 0 < ζ < 1 represents 15 an underdamped, gradually fading oscillation; ζ = 1 indicates critical damping (rapid exponential decay without oscillation); and ζ > 1 corresponds to overdamping (slower exponential decay).

The relaxation of slow-closure pores was less damped than that of kiss-and-run pores (median ζ = 1.13, range 0.79–1.88, versus 1.79, 0.98–3.63; n = 8 each; Mann–Whitney U test, p = 0.028; Figure 6A, D). Three slow-closure pores were underdamped (ζ = 0.79–0.96), and most others were near critical damping. By contrast, seven of eight kiss-and-run pores were overdamped. The fits to velocity had R² = 0.922–1.000 for slow-closure pores and 0.996–1.000 for kiss-and-run pores. The analysis did not produce underdamping, since when applied to pore trajectories simulated without oscillatory components it yielded only overdamped values (ζ = 1.15–2.16; Figure 6D). However, values of ζ close to 1 cannot distinguish a weakly underdamped oscillation from a two-phase monotonic closure.

**Figure 6.**
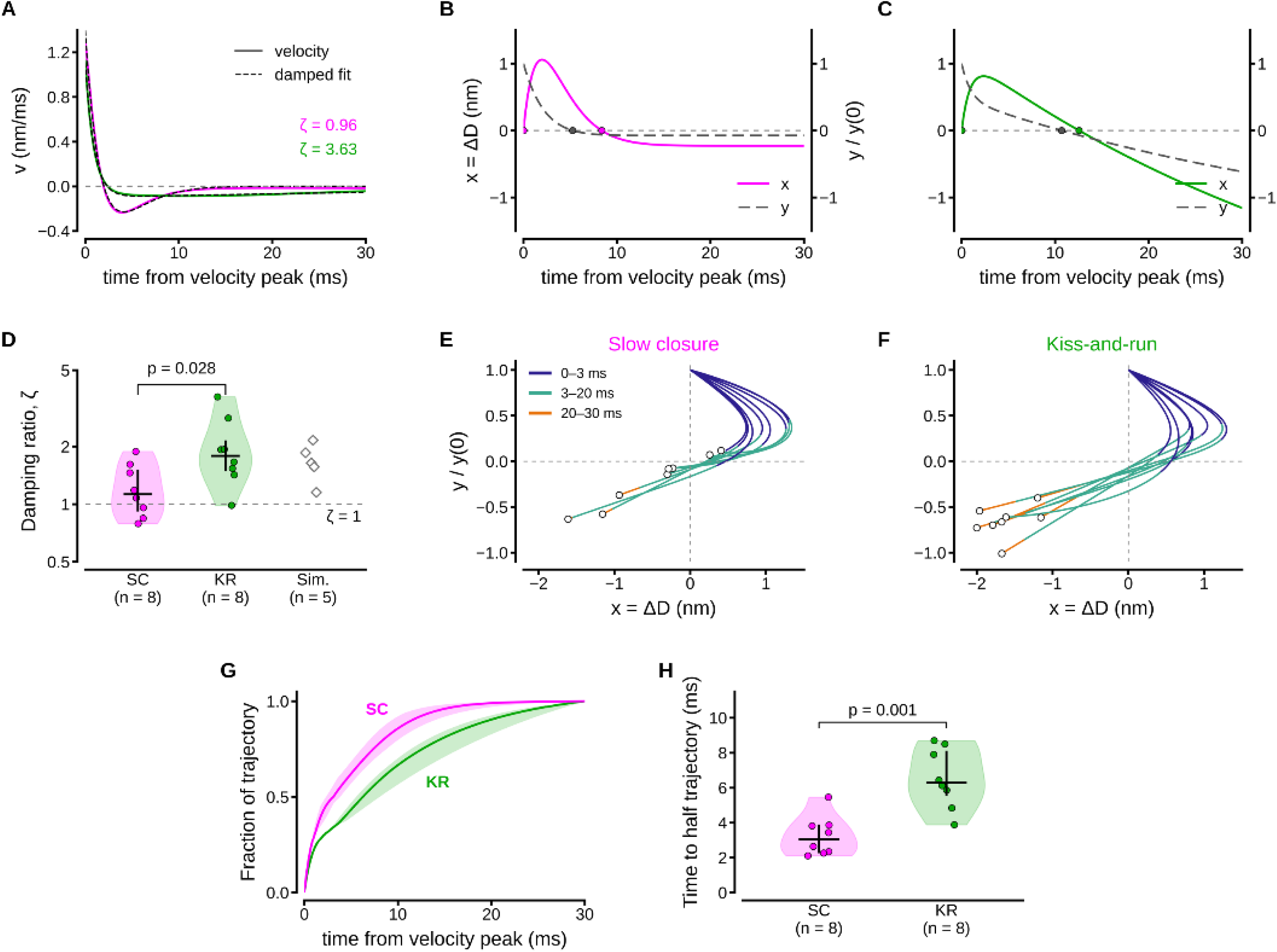
Damped relaxation of slow-closure and kiss-and-run pores. A) Pore velocity after peak of representative events (maximal velocity above 1 nm reached 0.5 ms after the pore crossed 1 nm). Dashed traces are fits to a damped second-order system. B, C) Displacement of pore diameter from its velocity peak value (x = ΔD, solid, left axis) and coupled internal process (y, dashed, right axis, normalized to the velocity peak) for slow-closure (B) and kiss-and-run (C) events; circles mark zero crossings. D) Damping ratio (log scale) for slow-closure (n = 8) and kiss-and-run (n = 8) events. Open diamonds are simulated pore trajectories without oscillatory components. E, F) Phase portraits of slow-closure (E) and kiss-and-run (F) events (same as in D). Colors represent time after the velocity peak; open circles mark its end at 30 ms. G) Fraction of the trajectory in the x–y plane completed over time (medians and interquartile ranges). H) Time to complete half of the trajectory; Mann–Whitney U test.

The second-order behavior in the undershoot of slow-closure pores implies a second variable coupled to the pore diameter. Because the second-order fit could not be interpreted in terms of mass and friction, it was instead expressed as the mathematically equivalent system of two coupled first-order equations, ẋ = −ax + by and ẏ = cx − dy. The term x is the displacement of the pore diameter from its value at the velocity peak, ΔD(t) = d(t) − d(t_peak_), obtained by integrating v(t) from x(0) = 0; the term y is a second state variable, referred to here as the force applied to the pore (Materials and methods). Physically, x can be interpreted as the pore diameter, and y as an external force acting on it. The pore moves against viscous friction under its own elastic restoring force and the external force, which builds as the diameter changes and relaxes with its own time constant. In this description, ζ is set by the relaxation rates of the pore and of the force relative to the strength of their coupling. Strong coupling or slow relaxation of the force lowers ζ and speeds closure, as in kiss-and-run pores. The opposite favors slow closure of pores.

In six of eight slow-closure pores and in the eight kiss-and-run pores, x fell below zero at 4.1–14.0 ms and 4.3–14.7 ms, respectively, as the pores closed (Figure 6E-F). In the other two slow-closure pores, x and y settled at positive values (Figure 6E). The phase portraits of the two pore types (Figure 6E, F) differed in the speed at which each pore moved along its trajectory. Slow-closure pores covered half of their trajectory within 3.0 ms (median; range 2.1–5.5 ms) after the velocity peak. This is about twice as fast as kiss-and-run pores (6.3 ms, 3.9–8.7 ms; n = 8 each; Mann–Whitney U test, p = 0.001; Figure 6G, H). Moreover, slow-closure pores completed 86% by 10 ms, compared with 67% for kiss-and-run pores (p = 0.005). The difference did not depend on the relative scaling of x and y, since, when measured along x alone, half of the trajectory was covered in 3.6 ms versus 7.1 ms (p < 0.001). Thus, slow-closure pores reached the end of their relaxation sooner, consistent with their faster undershoot shown in Figure 5D–F.

## Discussion

To our knowledge, this is the first evidence of quantal release through the fusion pore. Slow-closure events resemble the “kiss-and-stay” events of PC12 cells, small, long-lasting currents that accounted for about 10% of exocytotic events (van Kempen et al., 2011). However, slow-closure pores release quanta via different pore kinetics. It remains to be determined whether the two cases represent the same mechanism. One concern with our data is that applying force to the membrane to stimulate exocytosis could produce an artifactually slow release. However, membrane tension opens transmembrane ion channels and induces a normal pattern of vesicle fusion (Shi et al., 2018). Moreover, tension applied to the plasma membrane may induce complete vesicle fusion without affecting pore formation in vesicles that are already docked (Gao et al., 2008; Shendrik et al., 2023).

Another concern is the interpretation of Equation 1, our empirical description of the current. Its exponential terms do not by themselves distinguish between pores formed by the reorganization of large lipid populations and pores formed by transitions in a few proteins. The rise term has the form of n independent first-order steps, which could equally reflect a small number of protein units. Exponential kinetics are nonetheless consistent with lipid-driven pore formation (Chanturiya et al., 1997; Oleinick et al., 2013; Risselada et al., 2014; Kawamoto et al., 2015; De-Miguel, 2026a). Lipids take part in the structure of the stalk and the pore (Shinoda et al., 2010; Aeffner et al., 2012; Poojari et al., 2021). In simulations, lipid composition determines the rate at which the pore opens (Marrink and Mark, 2003), and membrane tension selects the fusion pathway and pore dynamics (Gao et al., 2008).

### The meaning of a quantum

Frozen pore extrapolation predicts that slow-closure and full fusion vesicles contain more molecules than are released (Figure 1D; De-Miguel, 2026a). This observation is consistent with the retention of part of the vesicle content in other cell types (Omiatek et al., 2010; Li et al., 2015; Ren et al., 2016). A quantum is therefore better defined as the amount released by one exocytotic event than as the content of a vesicle. In any case, release through a fusion pore is not fixed by the vesicle but set by the forces that keep the pore open.

### What the rise of the current reports

The delay at the onset of the current, captured here by the exponent n and the time constant τ_0_, reflects both the opening of the pore and the time required to hydrate the matrix and free the transmitter. Transmitter efflux is limited by the conductance of the pore (Breckenridge and Almers, 1987a; Alvarez de Toledo et al., 1993; Albillos et al., 1997; Lindau and Alvarez de Toledo, 2003), so an increase in current reports pore widening and the forces that drive it (Amatore et al., 2010; Oleinick et al., 2013). Differences in n and τ_0_ among release modes arose during the early, sub-nanometer stage, when hydration and pore nucleation overlap; once pores exceeded 1 nm, all three types expanded at similar rates. Hydration is unlikely to explain the differences between modes, because all events came from the same population of dense-core vesicles, with contents of the same order. Matrix hydration should therefore introduce a similar delay across modes. The fastest onset in feet preceding full fusion, would then reflect hydration plus larger or faster nucleating forces. On the contrary, the more delayed onset of slow-closure and kiss-and-run currents may indicate smaller or slower nucleating forces that fail to drive full fusion and leave the pore susceptible to closure.

### Closing dyamics

A surprising result is that slow-closure pores initially close faster than kiss-and-run pores, then settle at a residual diameter, and relax with less damping. Their longer onset delay suggests weak initial forces. Yet their current amplitude, pore area and early release exceed those of kiss-and-run pores. The energy stored while the pore expands cannot drive vibrations because inertia is negligible at this scale. It instead dissipates through viscous processes coupled to the pore diameter, such as relaxation of membrane tension or disassembly of the fusion complex (Montana et al., 2009). Their interplay with the pore can produce fast closure to a residual diameter. A purely lipidic pore of 1.2–1.9 nm is unlikely to be stable. With a line tension of 3–4 × 10⁻¹¹ N (Tolpekina et al., 2004a; Leontiadou et al., 2004; Wohlert et al., 2006), it would need a membrane tension of 35–60 mN/m. Simulated bilayers rupture near that range (Leontiadou et al., 2004), and small hydrophilic pores become unstable above about 4–5 mN/m (An, 2023). Finite-size pores can instead be stabilized by membrane fluctuations at low tension (Farago and Santangelo, 2005) or by the fusion machinery (Long et al., 2012; Oleinick et al., 2017). Long-lived pores narrower than 1 nm do occur in chromaffin dense-core vesicles, where slow release events last seconds (Zhang et al., 2023).

### Coupled hypothesis of pore expansion

The coupled description that emerges from the damped relaxation implies that closing kinetics are set by three quantities: 1. the rate at which the pore relaxes (its elastic stiffness divided by the friction of lipids and water); 2. the rate at which the force relaxes (1/τF), and 3. the strength of the coupling between them. Slow-closure pores, which were less damped, reached the velocity undershoot nearly twice as fast as kiss-and-run pores, consistent with stronger coupling relative to relaxation. A passive viscoelastic force may account for the two-phase closure and the velocity undershoot, but not for the underdamping found in three slow-closure pores. In the absence of inertia, passive elements relax with real time constants. Underdamping therefore requires a non-reciprocal coupling between pore and force, which implies a force fed by an energy source, such as active feedback that responds to changes in diameter with a delay.

### Continuity between early and late events

The notion that the early stage of the pore shapes its fate is supported by correlations between the rise of the current and the closure of the pore. Independently of pore type, τ_0_ was negatively correlated with ζ, since pores that rose with a longer delay closed with less damping. Likewise, the time to the current peak predicted the time to the velocity undershoot. By contrast, the rise exponent n did not correlate with any closing parameter, so it is the timing of the rise, not its shape, that anticipates the closure of the pore.

These early differences carry through to closure. Slow-closure pores closed in two phases, whereas kiss-and-run pores closed in a single phase. This behavior is consistent with a metastable intermediate state, a shallow energy minimum at the residual diameter that slow-closure pores reach but kiss-and-run pores do not. Since a lipidic pore of this size is unlikely to be stable alone, such a minimum requires support from the fusion machinery. Whether the residual diameter corresponds to a distinct energy minimum and how high the barrier to final closure is remain to be tested.

### Differences and similitudes of pore populations

Kiss-and-run and slow-closure pores form a single continuous population. Above 1 nm they open at the same rate, and their rise kinetics vary continuously between types. Differences in their dynamics are therefore more likely to arise from forces acting on the pore than from the pore itself. Consistent with this view, kiss-and-run and full fusion share a common molecular machinery (Alés et al., 1999; Lindau and Alvarez de Toledo, 2003). Some studies describe the modes as distinct. Amperometric spikes of neuroendocrine cells fall into three clusters (van Kempen et al., 2011), and events of dense-core vesicles in chromaffin cells fall into fast and slow classes with different pore properties (Zhang et al., 2023). The pore may also bifurcate after opening into full fusion or kiss-and-run, depending on vesicle size (Chang et al., 2022). Our data do not exclude such a bifurcation but place it after a common early stage whose kinetics vary continuously and correlate with the timing of closure.

### Possible origin of the forces

If the differences between modes arise from forces acting on the pore, the variable y in the coupled equations can be interpreted as the force exerted on the pore by its surroundings. This force is driven by the pore diameter (term cx) and acts back on the pore (term by), as expected for a force-generating process whose load changes as the pore widens or narrows. Slow-closure and kiss-and-run pores would then differ mainly in how strongly this force is coupled to the pore and how fast it relaxes. In the three underdamped slow-closure pores, the couplings have opposite signs (bc < 0); as discussed above, this non-reciprocal exchange requires an energy source.

The data do not identify this force, but three candidates stand out. The first is the fusion machinery at the rim of the pore. Together with lipids, the SNARE complex and synaptotagmin provide the force that opens and dilates the pore (Han et al., 2004; Han and Jackson, 2005; Lynch et al., 2008; Wu et al., 2021; Zhang et al., 2022). Movement of the SNARE transmembrane domains suffices to open it (Fang and Lindau, 2014; Sharma and Lindau, 2018), and crowding of SNARE complexes around it generates entropic forces that drive expansion (McDargh et al., 2018). The number of complexes matters. One suffices for liposome fusion 22 (van den Bogaart et al., 2010), two for small synaptic vesicles (Sinha et al., 2011), and at least three are needed for fast fusion of dense-core vesicles (Hua and Scheller, 2001; Mohrmann et al., 2010). More complexes produce faster and larger pores (Bao et al., 2018). The longer initial delay of slow-closure currents may therefore reflect fewer active complexes.

The second candidate is membrane tension, the small net result of large opposing lipid forces (Evans and Rawicz, 1990; Lindahl and Edholm, 2000), which allows partial release when cholesterol-rich membranes resist pore re-closure (Najafinobar et al., 2016). The third is the cytoskeleton. The actin cortex buffers membrane tension (Raucher and Sheetz, 1999; Neco et al., 2008; Doreian et al., 2008; Bretou et al., 2014; Trouillon and Ewing, 2014; Wen et al., 2016), branched actin expands and stabilizes the pore, and myosin II constricts it (Olivares et al., 2014; Biton et al., 2023). SNARE zippering, actin remodeling and myosin II contraction draw on stored or metabolic energy and could supply the active component required for underdamping. The available data cannot assign the force to any one of them.

### Possible calcium regulation of slow-closure

In the extrasynaptic exocytosis of the neurons studied here, as in endocrine cells, dense-core vesicles are not tethered to voltage-gated calcium channels as synaptic vesicles are. Fusion is driven by calcium released from intracellular stores (Trueta et al., 2004), in a feedback loop activated by the released serotonin (Leon-Pinzon et al., 2014). Intracellular stores also supply calcium in other forms of extrasynaptic release (Bao et al., 2003; Ludwig and Leng, 2006; Patel et al., 2009). Because cytoplasmic calcium diffuses poorly, vesicles at the center of a cluster may receive less calcium, owing to their distance from release sites and to surrounding vesicles acting as a barrier. The number of SNARE complexes determines whether a pore opens, stays open and expands (Mohrmann et al., 2010; Shi et al., 2012; Bao et al., 2018), and calcium can shift the mode of exocytosis, although in opposite directions in different preparations (Alés et al., 1999; Elhamdani et al., 2006). We therefore hypothesize that low calcium engages fewer SNARE complexes, reducing the expanding force and increasing the proportion of kiss-and-run or slow-closure events.

## Materials and methods

### Isolation and culture of neurons

Experiments were conducted using serotonergic Retzius neurons isolated from leeches (*Hirudo verbana*). Individual neurons were isolated from the ganglion one by one using a glass pipette, then sterilized and plated in culture (Dietzel et al., 1986). Experiments were performed at 18°C after 1-8 days in culture.

### Amperometric recordings, selection tracing and processing

Amperometric currents upon 5-HT release were recorded using conventional amperometry. Only events exceeding the average baseline noise by more than five standard deviations, without contamination from other spikes, were included in our analysis. Slow-closure events had a fast rise and biexponential decay, with amplitudes <10 nA. The kinetics of selected individual amperometric currents were noise-filtered using the “Statistical Empirical Mode Decomposition” (SMED) algorithm, which incorporates a signal-smoothing procedure (Kim et al., 2012).

### Accuracy of amperometric recordings

A serious concern in interpreting current kinetics is ensuring that the measured current reflects release at the pore rather than diffusion over long distances between the pore mouth and the carbon fiber. We estimated the maximum distance compatible with undistorted kinetics as the root-mean-square displacement of 5-HT, √(2Dt), during the rise time of the recording system, t ≈ 34 µs (0.34/fc for the 10-kHz Bessel filter). Using the free-solution diffusion coefficient of 5-HT (5.9 × 10⁻⁶ cm²/s, from the Stokes–Einstein relation), this distance is 200 nm. If diffusion between the membrane and the fiber were as 24 hindered as within the pore, using the common diffusion coefficient for the reconstructions (D = 4.49 × 10⁻⁷ cm²/s; De-Miguel, 2026a), the distance would be 55 nm. The pressure applied with the carbon fiber to stimulate exocytosis kept the fiber in contact with the membrane and may have helped keep the release sites within this range. Larger gaps mainly slow and smooth the rising phase, which is also influenced by the slow release of 5-HT as the intravesicular gel hydrates and by the small amount of transmitter leaking through the nascent narrow pore (Hafez et al., 2005). By contrast, the peak and the decay, which last tens of milliseconds, are hardly affected. Under these conditions, the current reflects the dynamics of the pore and the amount of 5-HT released, except during the early rising phase.

### Empirical model of the kinetics of 5-HT release

Amperometric currents containing individual opening and closing events were fitted to the empirical equation:

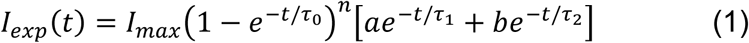

where *I_max_* is the maximum current amplitude. The first factor in Equation 1 describes the rise of the current; the second factor, represented by the sum of two decaying exponentials, describes the current decay produced by slow-closure pores. The exponent *n* in the rising phase provided the best fit to the rise of currents. Fitting the filtered currents to Equation 1 generated the amplitude parameters a and b, the rise time constant *τ*_0_, and the decay time constants *τ*_1_ and *τ*_2_. Slow-closure currents had *τ*_1_ and *τ*_2_ values that differed by about one log unit. The *R*^2^ oeffiiiintt iterpretedd ss hee tandardd eviitiinn ff hee esiiualss determined the quality of the fits

For completeness, we also fitted the current kinetics to a stretched-exponential form, which may explain pore formation as crack-like behavior of the plasma membrane (Brandt and Edholm, 2010). However, this fit was less accurate (R^2^ < 0.86, compared with 0.98 for Equation 1). Alternatively, we used a sigmoidal equation to describe the rising phase of the amperometric currents, as it provides a cooperative kinetic interpretation. However, these fits were also of lower quality than those obtained with Equation 1, as evidenced by a lower correlation coefficient and a wider range of root-mean-square values (0.31 to 0.86) for the sigmoidal equation, compared with 0.02-0.17 for Equation 1.

### Toroidal pore reconstruction and frozen-pore currents

Pore reconstructions used a toroidal pore geometry and the same diffusion coefficient as for the kiss-and-run events in De-Miguel (2026a). In brief, the pore was modeled as the neck of a torus with curvature radius ρ_c_ = 7.37 nm, where the minimal radius r is the pore radius. The 5-HT diffusion coefficient was D = 4.49 × 10^−7^ cm^2^/s, the vesicle radius was 51.85 nm, and the charge per oxidized molecule was 4e.

### Pore diameter at the current peak

The diameter at the current peak was the hydrated flux packing (HFP) diameter (De-Miguel, 2026a), d_HFP_ = d_H_(R/r)_N_, where d_H_ = 0.85 nm is the hydrated diameter of 5-HT, N is the peak flux in molecules per microsecond rounded up to the next integer, and (R/r)_N_ is the ratio between the radius of the smallest circle that encloses N equal circles and the radius of those circles.

### Vesicular content

The 5-HT content of each kiss-and-run and slow-closure vesicle was estimated by frozen pore extrapolation, FPE, as described in De-Miguel (2026a). In brief, N_FPE_ = q(t_peak_) + J_peak_τ, where q(t_peak_) is the number of molecules released up to the current peak, J_peak_ = I_peak_/4e is the peak flux and τ = V_ves_L_eff_/(Dπr^2^) is the emptying time constant of a toroidal pore of diameter d_HFP_. For the foot of full-fusion currents, the mean quantum of full-fusion events (91,585 molecules) was used; its charge (58.7 fC) is close to that of somatic dense-core-vesicle events of Retzius neurons (47–52 fC; Bruns et al., 2000).

### Frozen-pore currents

The current that a pore frozen at time t_0_ would produce (De-Miguel, 2026a) is I_fr_(t) = 4e[N_FPE_ − q(t_0_)]/τ_fr_ × exp[−(t − τ_0_)/τ_fr_], with τ_fr_ = V_ves_L_eff_(r_0_)/(Dπr_0_^2^), where r_0_ is the pore radius at τ_0_; by construction, I_fr_(τ_0_) equals the recorded current. Pores were frozen at the current peak (r_0_ = d_HFP_/2) or at the transition between the fast and slow decay components of slow-closure currents.

### Model-free analysis of currents

Recorded currents were quantified by time to peak, half-width, fraction of charge released before the peak, number of molecules released within 16 ms (the duration of the shortest recorded current, 16.3 ms), and number released between 16 ms and the end of the recording. The decay from the peak to 16 ms was fitted with one and two exponentials, using time constants from a logarithmic grid of 28 values between 0.3 and 60 ms and amplitudes obtained by non-negative least squares. The fits were compared using the Bayesian information criterion (BIC); a difference above 10 was taken as strong evidence for two exponentials. Groups were compared with exact two-sided Mann–Whitney U tests (Mann and Whitney, 1947) without correction for multiple comparisons; the comparisons tested two hypotheses stated in advance, that the early phase is the same and that release after the early phase differs.

### Reconstruction of the dynamics of pore diameter

The diffusional resistance of the pore, R(t) = [N_FPE_ − q(t)]/[J(t)V_ves_], was converted to a diameter by solving L_eff_(r)/(πr^2^) = DR(t) for r. By construction, the diameter at the current peak equals d_HFP_. Reconstructions ended when less than 5% of the vesicular content remained. Segments computed from the exponential extrapolation of the fitted currents beyond the end of the recording are shown as dotted lines. Pore velocity was calculated as the first derivative of a cubic Savitzky–Golay filter applied to the reconstructed diameter (Savitzky and Golay, 1964).

### Opening and closing phases of the pore

Within the reconstructed range, the pore velocity peaked at the first analyzed point, 0.5 ms after the pore crossed 1 nm (half the filter window is lost at the edge); the absolute velocity peak occurred below 1 nm (0.4–1.1 nm) in all events. This maximal velocity is referred to as the velocity peak (Figure 6). To time the transitions between opening and closing, the diameter from the crossing of 1 nm was fitted over 30 ms with the sum of two exponentials, d(t) = d∞ + A_1_exp(−t/τa) + A_2_exp(−t/τb), describing two coupled first-order relaxations without inertia (R² = 0.987–1.000). The end of opening was defined as the peak of the fitted diameter; closure was defined as the time at which 95% of the decline from the peak to d_∞_ had been completed. The closing phase was fitted from the time of maximal closing velocity with one or two decaying exponentials, d(t) = d_∞_ + B_1_exp(−t/τ_fast_) + B_2_exp(−t/τ_slow_); the two-exponential model was retained when it lowered the Akaike information criterion by more than 10 (Akaike, 1974). The transition between the fast and slow closing phases was defined as the time at which both phases had equal closing velocities, t = ln[(B_1_/τ_fast_)/(B_2_/τ_slow_)]/(1/τ_fast_ − 1/τ_slow_). The closing time constant was that of the fast phase in slow-closure pores. For the slow-closure event of Figure 5C, the fits were restricted to the interval before its minimal diameter (11.3 ms). The velocity undershoot was measured from the diameter velocity after crossing 1 nm, the end of opening being zero crossing; the time to the undershoot was the time from the end of opening to the velocity minimum; the half-width was the time between the crossings of half that minimum before and after it; the recovery was the time from the minimum to the later crossing.

### Damped second-order model

Velocity over time was fitted to the solution of a damped second-order linear system, ṽ + 2ζω_n_v̇ + ω_n_^2^v = 0. The model has three conditions: underdamped (ζ < 1), critically damped (ζ = 1), or overdamped (ζ > 1). Fittings were made using nonlinear least squares with the Levenberg–Marquardt algorithm (Levenberg, 1944; Marquardt, 1963). The relaxation was characterized by the damping ratio ζ and the natural angular frequency ω_n_. In underdamped conditions, the oscillating velocity decays with a time constant of 1/(ζω_n_); in critically damped and overdamped conditions, the velocity returns to zero without oscillation, as a sum of two exponentials when overdamped. Of the 10 slow-closure and 10 kiss-and-run events, two slow-closure events were excluded: the event in Figure 5C, whose pore re-expanded late in the reconstruction, and one of the three events whose fits reached the upper bound of ζ (50); the other two were kiss-and-run events whose pores closed as a broad hump (Figure 4—figure supplement 1A) Fits with ζ fixed at any value above about 5 were equally good (R² changed by less than 0.002), so their damping ratio was undetermined.

### Estimates of forces without inertia

At the scale of the fusion pore, the low Reynolds number implies that viscous forces far exceed inertial forces. The pore diameter therefore obeys a first-order overdamped equation, γẋ = −kx + F(t). Here γẋ is the viscous drag, −kx the elastic force restoring the pore to equilibrium, and F(t) any external force. With a constant force, the pore relaxes monotonically. Second-order behavior arises when the force is dynamic. If the force builds in opposition to the change in diameter and relaxes with a time constant τF, dF/dt = −κx − F/τF, and the two equations form a coupled system with a = k/γ, b = 1/γ, c = −κ and d = 1/τF. Here γ is the friction coefficient of the pore (viscous drag of lipids and water), k the elastic stiffness of the pore neck (line tension and bending rigidity), κ the rate at which a change in diameter builds the force, and τF the relaxation time of the force. Then ω_n_² = ad– bc = k/(γτF) + κ/γ and 2ζω_n_ = a + d = k/γ + 1/τF, where ω_n_ is the natural angular frequency, the angular frequency of the system without damping. Velocity obeys the same equation, ṽ + 2ζ ω_n_ v̇ + ω_n_² v = 0. The eigenvalues, λ = −β ± I ω with β = ζω_n_ and ω = ω_n_(1 − ζ²)^1/2^, are complex when (a − d)² + 4bc < 0, that is, when κ > γ(k/γ − 1/τF)²/4. Underdamping thus requires a force that opposes the displacement of the pore, not only its velocity. Then x and y cross zero out of phase and the trajectory spirals toward the origin; with real eigenvalues, both decay without oscillation (Strogatz, 2015). A passive viscoelastic element, whose force responds to velocity (dF/dt = −κv − F/τF), yields only real eigenvalues. Without inertia, a passive linear system that only dissipates energy relaxes without oscillation, whatever its number of internal variables, although it can produce a two-phase closure, a velocity undershoot and a single crossing of the final diameter. The fit determines only ω_n_ and ζ, so γ, k, κ and τF cannot be identified separately.

To plot trajectories, we used the symmetric decomposition a = d = ζω_n_, with b = −c = ω_n_ (1 − ζ²)^1/2^ for ζ < 1 and b = c = ω_n_ (ζ² − 1)^1/2^ for ζ > 1. This convention is not unique and does not give the physical coefficients. Then y = (v + ζω_n_ x)/b. x is expressed in nm, and y was normalized to its value at the velocity peak, y/y(0) = (v + ζω_n_x)/v(0), which is independent of b and leaves the zero crossings unchanged. Progress along the trajectory was measured as the cumulative path length in the x–y plane, normalized to the path length at 30 ms after the velocity peak. The time to half trajectory was the time at which this fraction reached 0.5. To test the dependence on the relative scaling of x and y, the same measure was computed along x alone.

### Robustness tests

To test whether the analysis produces underdamping, currents were simulated without oscillatory components (single-exponential closures with time constants of 10 and 20 ms, or two-phase closures with a fast component of 3–5 ms followed by a slow component of 40–300 ms). We used the toroidal geometry and the same diffusion coefficient. Data were analyzed with the same reconstruction and fit, using either the true or the FPE vesicular content (Figure 6D). The dependence of the damping ratio on the fit window was evaluated using windows of 15, 25, 30, and 40 ms. To test the reconstructions’ sensitivity to the vesicular content, reconstructions were repeated with the FPE content multiplied by 0.90, 0.95, 1.05, and 1.10. Damping ratios changed little, whereas diameters near vesicle depletion changed markedly.

### Correlations between rise and closure

Spearman correlations were computed between four rise parameters (τ_0_ and n of Equation 1, time to peak, maximal rate of rise) and five closure parameters (ζ, ω_n_, time to undershoot, undershoot half-width, closing time constant) for the 16 pores of the damping analysis, pooled and within pore types. Within types, values were ranked separately in each type and significance was assessed with 5,000 within-type permutations, so that differences between types could not produce a correlation. The τ_0_–ζ correlation was hypothesized from earlier analyses; the other 19 were exploratory. After Holm correction for 20 comparisons only the within-type correlation between time to peak and time to undershoot remained significant (p = 0.02).

## Acknowledgements

Data used in this paper were obtained by Montserrat Guerrero for different purposes in the laboratory of the corresponding author. We wish to express our gratitude to Sara Flores González for her excellent assistance with animal and laboratory care. We also wish to acknowledge Mr. Francisco Pérez and Antonio Torres Galván from the computer unit of IFC-UNAM for his technical support.

## Author contributions

Alejandro Martínez-Valencia, Investigation, Formal analysis; Guillermo Ramírez-Santiago, Formal analysis, Methodology; Francisco F. De-Miguel, Conceptualization, Supervision, Funding acquisition, Formal analysis, Writing and editing

## Data availability

Amperometric recordings, fitted currents and the analysis code used to generate Figures 1–6 will be deposited in a public repository with a DOI, to be cited here and in the reference list.

## Ethics

Experiments were performed on cultured neurons isolated from leeches (Hirudo verbana), an invertebrate species not subject to regulations for vertebrate animal research; animals were anesthetized on ice before dissection.

## Supplementary Figures

**Figure 1—figure supplement 1.**
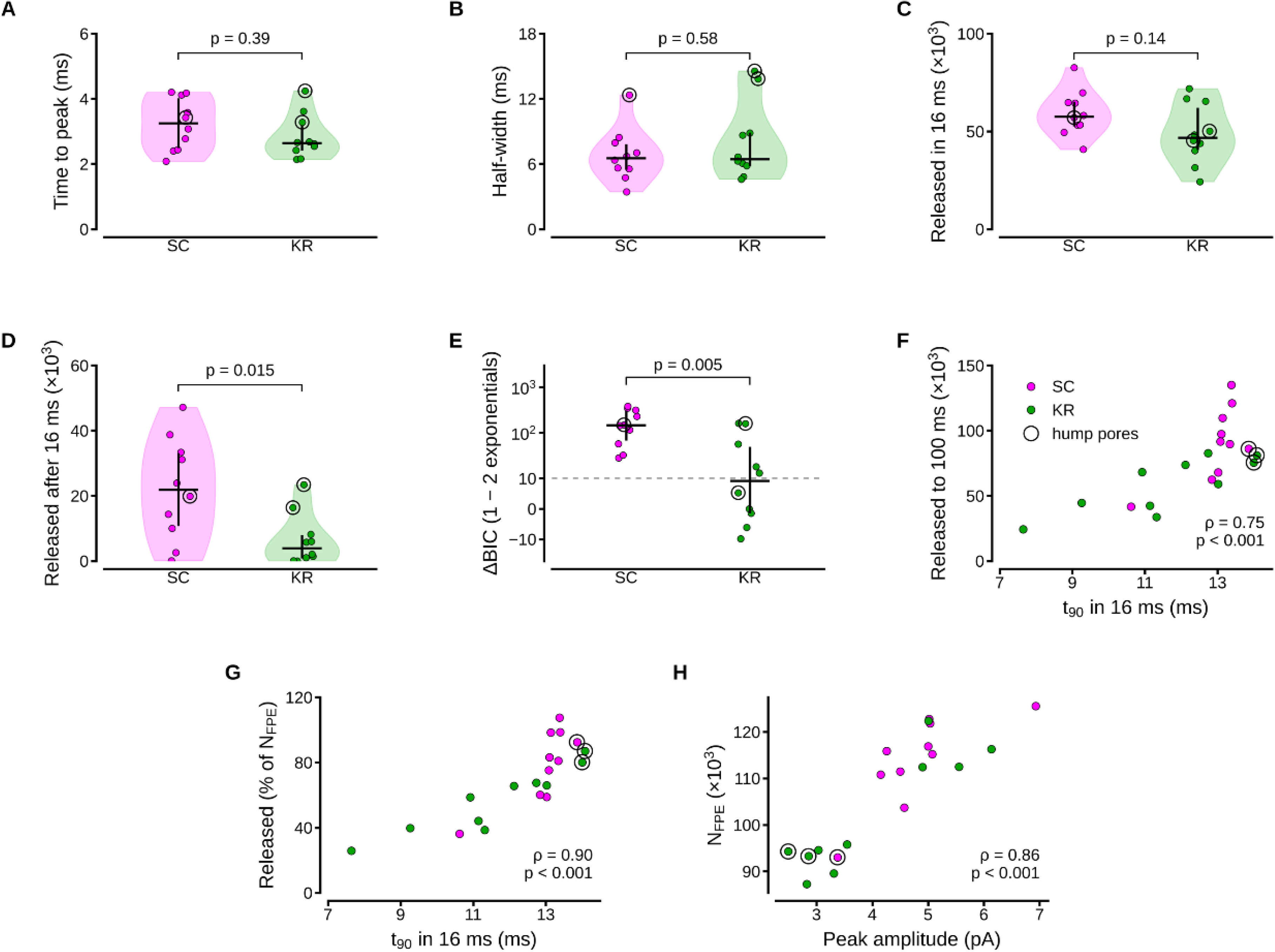
Model-free comparison of slow-closure and kiss-and-run currents. (A–C) Early phase. Time to the current peak (A), half-width (B) and number of 5-HT molecules released in the first 16 ms (C), the duration of the shortest recorded current. (D) Molecules released after 16 ms by recorded currents without extrapolation. (E) Difference in the Bayesian information criterion (ΔBIC) between single-and double-exponential fits of the current decay from peak to 16 ms; values above 10 (dashed line) favor two exponentials. (F) Molecules released up to 100 ms versus the time taken to release 90% of the charge of the first 16 ms (t_90_); the extrapolated tail contributed a median of 10% (slow closure) and 3% (kiss-and-run) of this number; Spearman correlation. (G) Percentage of N_FPE_ released versus t_90_. (H) N_FPE_ versus peak current amplitude. In F–H, Spearman correlations. Slow-closure events, magenta; kiss-and-run events, green; n = 10 each; circles mark the three pores with a broad hump (Figure 4—figure supplement 1A). Horizontal bars, medians; vertical bars, interquartile ranges; exact two-sided Mann–Whitney U test.

**Figure 4—figure supplement 1.**
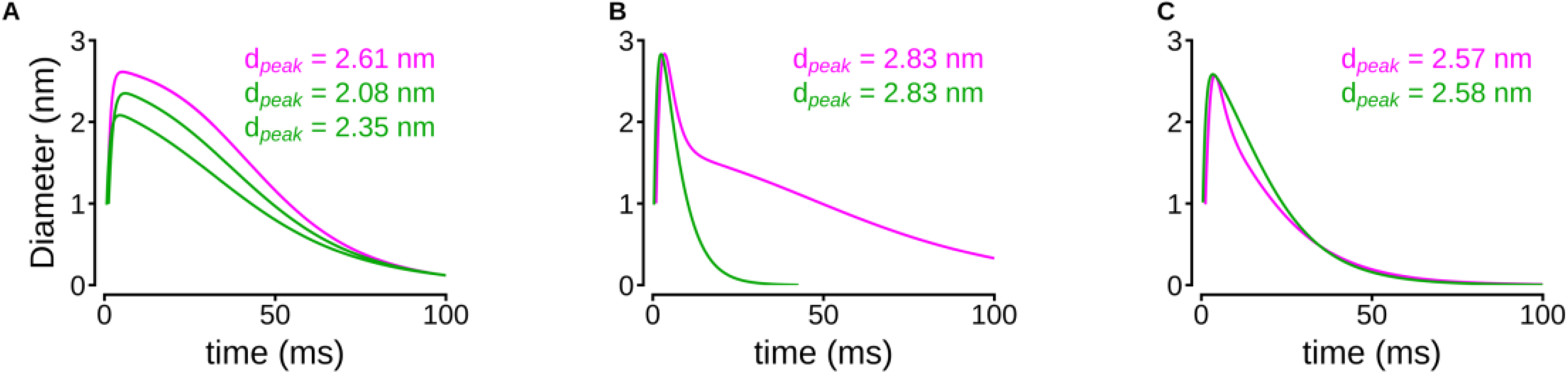
Pores with a broad hump and pores matched by peak diameter. A) The slow-closure pore (magenta) and the two kiss-and-run pores (green) that closed as a broad hump. B, C) Slow-closure and kiss-and-run pores matched by peak diameter (difference ≤ 0.05 nm). Peak diameters are indicated in each panel.

**Figure 5—figure supplement 1.**
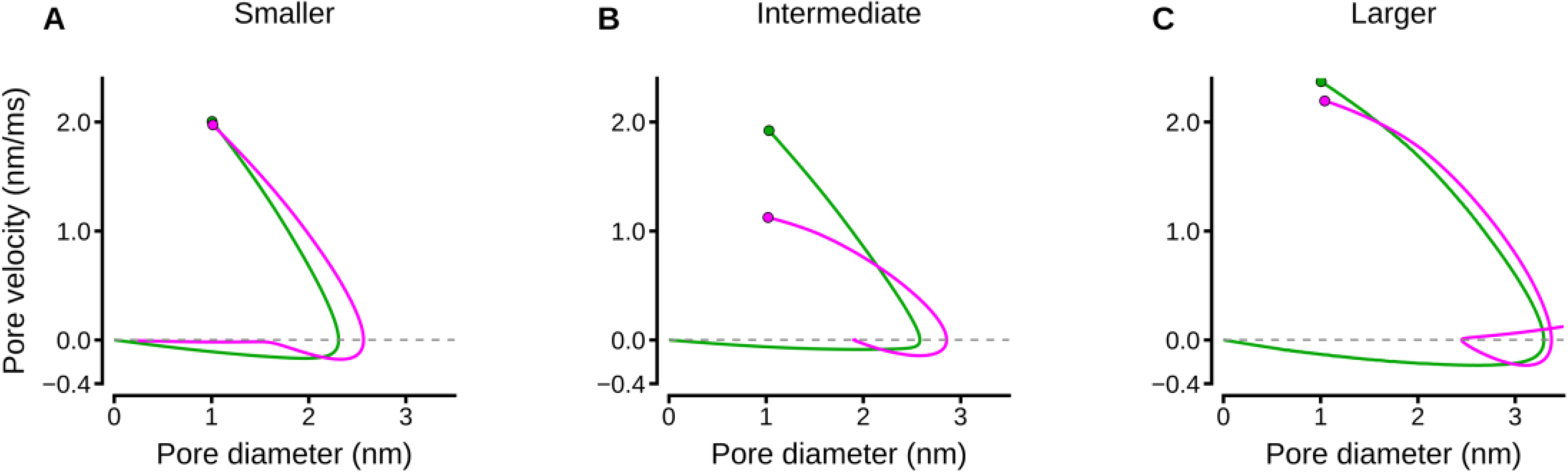
Pore velocity versus diameter. A-C) Trajectories of the pairs in Figure 5; circles mark the trajectory start at 1 nm. Part of each trace is extrapolated from the fitted current (Materials and methods).

**Figure 6—figure supplement 1.**
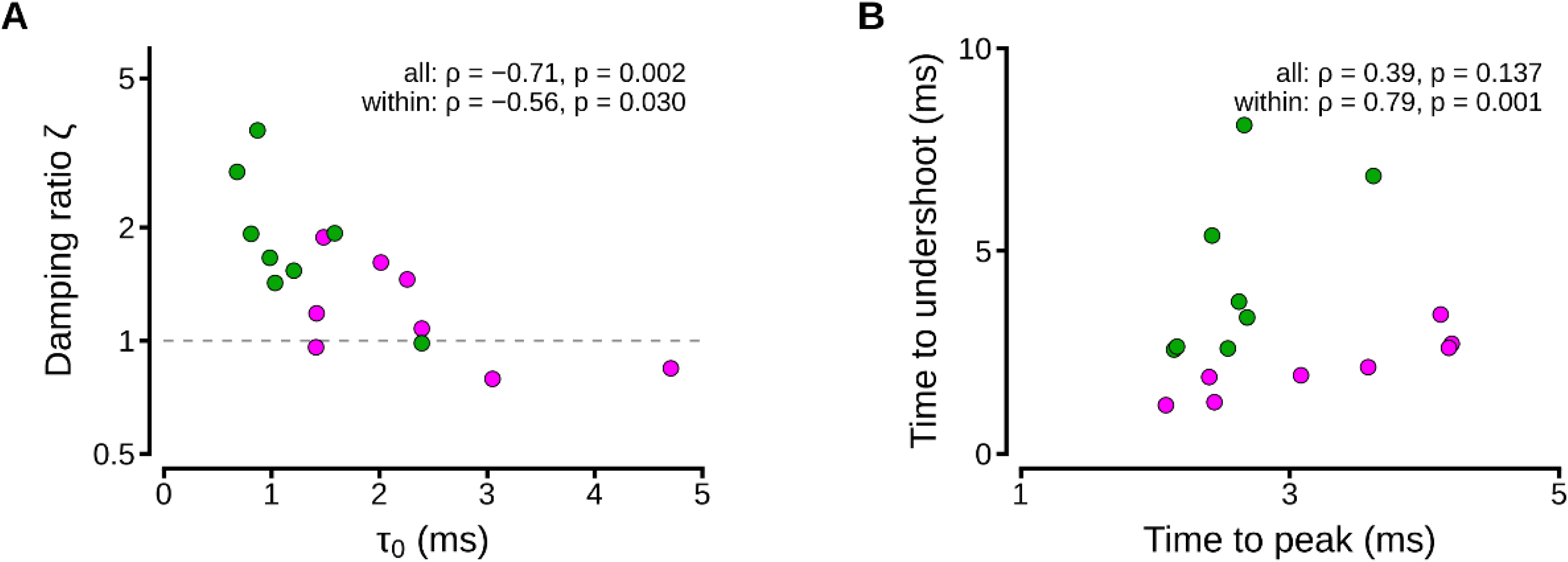
The rise of the current anticipates the closure of the pore. (A) Damping ratio ζ (logarithmic scale) versus the rise-time constant τ_0_ in Equation 1; the dashed line marks critical damping (ζ = 1). (B) Time from the end of opening to the velocity minimum (time to undershoot) versus the time to the current peak. Slow-closure (magenta) and kiss-and-run (green) pores from the damping analysis (n = 8 each). Spearman correlations are reported for all 16 pores and within pore types (ranks within each type; permutation tests stratified by pore type).

